# Engineering of Sr33 and Sr50 plant immune receptors to alter recognition specificity and autoactivity

**DOI:** 10.1101/2022.03.05.483131

**Authors:** Janina Tamborski, Kyungyong Seong, Furong Liu, Brian Staskawicz, Ksenia V Krasileva

**Affiliations:** Department of Plant and Microbial Biology, University of California Berkeley, USA; Innovative Genomics Institute, UC Berkeley, USA

## Abstract

Plants possess cytoplasmic immune receptors called nucleotide-binding leucine-rich repeat receptors (NLRs) that recognize the presence of a pathogen through a range of mechanisms: direct binding of effectors or indirect recognition of effector actions. The direct binding of effectors has been shown to be mediated through the NLR’s leucine-rich repeat (LRR) domain. Accurate prediction of amino acids involved in these direct interactions can greatly enhance understanding of effector recognition and inform efforts to engineer new resistance. In this study, we utilized two homologous NLR resistance genes from wheat, Sr33 and Sr50, that recognize distinct effectors by directly binding to them through their LRR domain. While the effector recognized by Sr50 is known and described as AvrSr50, the effector recognized by Sr33 remains unknown. Through a combination of phylogenetics, allele diversity analysis in the LRR and structural modeling, we identified the amino acids in Sr50 likely to physically interact with its effector. Mutation of these sites helped identify 12 amino acids we hypothesized to be sufficient to mediate effector binding in Sr50. Changing these 12 corresponding amino acids in Sr33, we showed AvrSr50-dependent initiation of cell death in wheat protoplasts and *Nicotiana benthamiana*. Furthermore, we were able to pinpoint and change amino acid residues that govern autoactivity of Sr50 in the wheat protoplast cell death assay. These findings are a major advance towards the successful engineering of new effector recognition specificities in direct binder NLRs.

## Introduction

Nucleotide-binding leucine-rich repeat receptors (NLRs) are intracellular immune receptors that recognize pathogen-derived molecules, called effectors, that are secreted into the plant cell by a pathogen with the goal of subverting plant immune responses. NLRs deploy different means of monitoring the cell by either directly binding to effectors or recognizing the action of an effector on a plant component. Plant components whose modification is recognized by NLRs are referred to as guardees or decoys. The recognition of an effector leads to the activation of downstream responses which culminate in localized cell death, referred to as hypersensitive response (HR) (Dodds and Rathjen, 2010). Although plant NLRs have been shown to evolve rapidly and recognize a wide range of structurally diverse effectors, robust approaches for rational modifications of NLRs and their recognition specificities remain limited (Tamborski and Krasileva, 2020).

Most NLRs share a common, multi-domain structure including a variable N-terminal domain, an NB-ARC domain, and a leucine-rich repeat (LRR) domain. There are three types of N-terminal domains that enable us to separate NLRs into three categories: CNLs that carry a coiled-coil (CC) domain, TNLs that carry a Toll/interleukin-1 receptor (TIR) domain and RNLs that have an RPW8-like domain (Shao et al., 2016). Furthermore, a subset of NLRs have one or more atypical plant protein domains integrated into their canonical NLR structure and are referred to as NLR-IDs (for integrated domain) (Grund et al., 2019). NLR-IDs usually function in pairs and are often genetically linked in a head-to-head orientation and are thought to exemplify the specialization of NLRs into a sensor and signaler function (Feehan et al., 2020). The integrated domains act as effector binding platforms and have been shown to directly interact with effectors (Cesari et al., 2013; Maqbool et al., 2015).

Several studies have already successfully engineered recognition specificities of NLRs. In direct binder NLRs, recognition was transferred from one NLR to another through chimeras (Ellis et al., 1999; Shen et al., 2003; Slootweg et al., 2017) and effector recognition was expanded through the means of random and targeted mutagenesis (Racman et al., 2001; Segretin et al., 2014; Chapman et al., 2014; Giannakopoulou et al., 2015). Also the modification of a guardee has proven successful through the introduction of effector substrate sites (Pottinger et al., 2020; DeYoung et al., 2012; Kim et al., 2016). Recently there have been some major advances in engineering the integrated domains of NLR-IDs to broaden or change their recognition specificity (Cesari et al., 2021; De la Concepcion et al., 2019; Liu et al., 2021). A major step forward was the identification and subsequent alteration of specific residues that mediate effector recognition. These successful engineering efforts exemplify that information from structural and effector binding studies help us not only understand but also engineer binding specificities of NLRs. While these most recent studies focused on modifying the integrated plant protein domain that binds to the effector, we have yet to make significant progress in engineering NLRs that bind to effectors through their LRR domain.

The recent cryo-EM structures of plant NLR genes have greatly enhanced our understanding of activated immune receptor function (Martin et al., 2020; Wang et al., 2019a, 2019b; Ma et al., 2020). The CNL ZAR1 and the kinase RKS1 guard the PBL2 kinase and initiate immune signaling upon its modification (Lewis et al., 2010; Wang et al., 2015). ZAR1 and RKS1 exist in pre-formed complexes that assemble into pentamers upon effector-mediated modification of PBL2 (Wang et al., 2019a, 2019b). The CC domains in this pentamer form a funnel-like structure that was shown to be a calcium-permeable channel (J. Wang, Hu, et al. 2019; Bi et al. 2021). Two groups independently defined the structures of the two direct binder TNLs RPP1 and ROQ1 and found that they assemble into tetramers upon activation through their cognate effectors. These studies confirmed the importance of intramolecular interactions to maintain the inactive state of NLRs and intermolecular interactions for resistosome assembly and immune signaling.

NLRs have provided a valuable source of disease resistance in many crop species. Wheat stem rust disease is caused by the fungal agent *Puccinia graminis* f.sp. *tritici* (*Pgt*) and has been a major threat to wheat production, especially since the emergence of the highly virulent *Pgt* race Ug99. A large number of resistance genes that protect against Ug99 have been cloned and deployed in recent years (Zhang et al., 2021; Saintenac et al., 2013; Chen et al., 2017; Steuernagel et al., 2016; Arora et al., 2019). Among them are Sr50 and Sr33: two CNL resistance genes that are highly similar with an amino acid identity of ~80% and provide resistance against wheat stem rust (Mago et al., 2015; Periyannan et al., 2013). They have been introgressed into wheat from *Secale cereale* and *Aegilops tauschii*, respectively, and are closely related homologs of the barley *MLA* (RGH1) family of NLR genes (Mago et al., 2015; Periyannan et al., 2013; Halterman et al., 2001; Wei et al., 2002). This family has been associated with high allelic and functional diversification, exemplified by the more than 30 barley MLA alleles that confer resistance to powdery mildew (Seeholzer et al., 2010).

Sr50 has been shown to directly bind to its cognate effector AvrSr50 (Chen et al., 2017). The effector recognized by Sr33 remains unknown to date, but is assumed to be distinct from AvrSr50 based on resistance to different *Pgt* races. (Mago et al., 2015). Sr50 and Sr33 are interesting candidates to investigate as their autoactivity seems directly linked to the level of expression as observed by (Cesari et al., 2016) and this study, which suggests that they might be in a sensitive on-off-state. Moreover, with 80% sequence identity the recognition specificities of these NLRs must be defined by a small number of amino acids in their LRR regions.

Here we report the engineering of Sr33 to recognize the effector AvrSr50. We used a combination of phylogenetics and structural modeling to predict the region within Sr50’s LRR that mediates binding to AvrSr50 and to identify individual residues involved in the interaction. We confirmed that the predicted region was sufficient to enable effector binding through the generation of a chimeric NLR that we named Sr33/50. This chimeric NLR was able to recognize and initiate cell death to AvrSr50 in wheat protoplasts and *Nicotiana benthamiana*. We subsequently generated the synthetic NLR Sr33_syn_ in which groups of residues within Sr33’s LRR were changed to the corresponding sites of Sr50. The exchange of 12 amino acids was sufficient for Sr33 to initiate cell death in wheat protoplasts and *N. benthamiana* in the presence of AvrSr50. We also observed that Sr50 could exhibit autoactivity in the wheat protoplast cell death assay, whereas Sr33 did not. We therefore used structural modeling to identify potential sites that could mediate this phenotype. We identified two residues that appear to be in close contact with the NB-ARC domain and mutation of these sites led to a stark reduction in autoactivity of Sr50. This study therefore demonstrates that the combination of sequence diversity analyses and structural modeling can be sufficient to identify and subsequently alter sites mediating effector recognition and autoactivity in direct binder NLRs.

## Results

We have previously identified highly variable NLRs (hvNLRs) by analyzing natural diversity of pangenomes of two model species, *Arabidopsis thaliana* and *Brachypodium distachyon* (Prigozhin and Krasileva, 2021). The highly variable amino acid residues within hvNLR homology groups together with structural modeling were predictive of the cognate effector-binding sites in *Arabidopsis thaliana* (Prigozhin and Krasileva, 2021). We decided to translate this knowledge to crop species and test if the same approach can accurately predict residues mediating NLR-effector interaction and subsequently guide rational engineering of NLRs. For this work we chose the wheat hvNLRs Sr50 and Sr33, that have been independently introgressed into bread wheat from *A. tauschii* and rye, respectively (Fig. 1B). We initially assessed sequence diversity of the Sr33 and Sr50 homology groups with homologs obtained with BLAST search against the non-redundant database in The National Center for Biotechnology Information (NCBI). The multiple sequence alignments (MSA) of each homologous groups indicated there was sufficient sequence diversity to predict binding residues in Sr50 (D. Prigozhin, personal communication; Krasileva, Prigozhin, Staskawicz, Liu (2021). Patent No. 113469, WO: U.S.). This approach proves to be an easily accessible way to obtain sequences for Shannon entropy calculation. However, as Sr33 and Sr50 displayed ~80% sequence identity, many collected homologs were shared by the two groups. We therefore needed to refine the initial analysis through phylogenetic analysis of the MLA family to curate Sr33 and Sr50 homologous groups with non-overlapping sets of sequences. We inferred the phylogenetic tree with MLA family members collected from Pooideae species, 14 wheat genomes (Table S1) and selected experimentally validated NLRs (Kourelis et al., 2021). Despite complex evolution within Triticeae species (Fig. 1B), we were able to resolve the NLRs into four distinct phylogenetic sub-families representing Sr50, Sr33, MLA1 and HcMLA1 grouped by similar phylogenetic distances. This allowed us to calculate Shannon entropy for each phylogenetic group with at least 13 unique NLR sequences (Fig. 1C; Table S2). The entropy plots of these four phylogenetic groups appear almost identical across the whole NLR length, suggesting that the same positions of variable or invariable sites are conserved across the MLA gene family.

**Fig 1.**
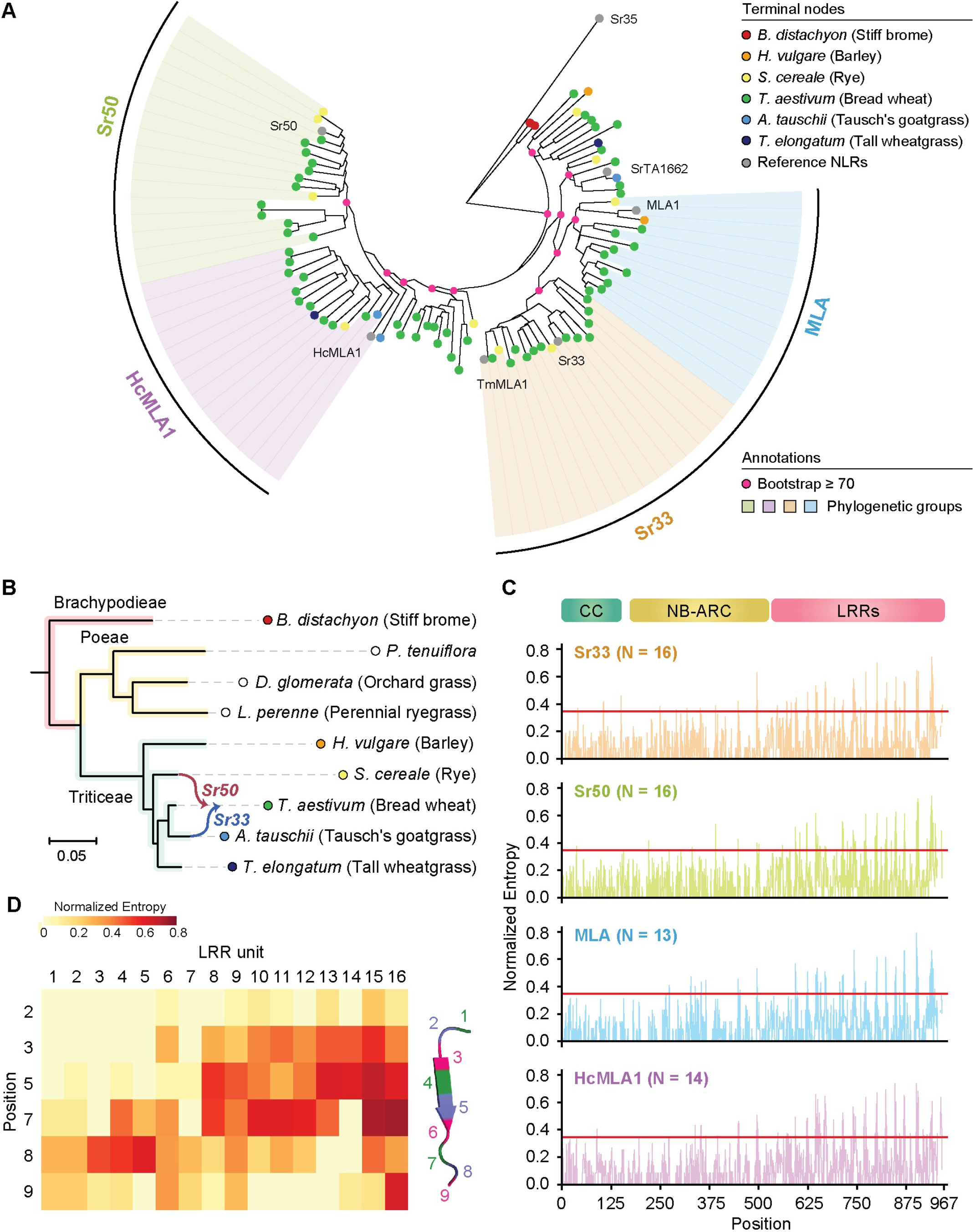
Evolutionary analyses and Shannon entropy scores of Sr33 and Sr50 related MLAs reveal variable hotspots in the NLR structure. **A**. The phylogenetic tree of selected MLA family members. The bootstraps are provided only for nodes used to distinguish the four phylogenetic groups or higher. The four phylogenetic clades were named based on the representative, experimentally validated NLRs that belong to the groups: Sr50, HcMLA1, Sr33 and MLA. All phylogenetic groups are supported with bootstrapping values ≥ 98, except for the HcMLA1 family that has a bootstrapping value of 63. The tree was rooted with the outgroup Sr35. **B.** The species tree of Pooideae members. The source of *Sr33* and *Sr50* introgression is indicated, and branches are colored based on tribe membership. No MLA family members were identified from the Poeae species, and these species are indicated with empty circles. **C**. The normalized Shannon entropy of the four phylogenetic group members in A. The normalized Shannon entropy was calculated with the indicated number of members (N) in each phylogenetic group. The normalized Shannon entropy ranges from 0 (no variability) to 1 (complete variability), and the red line indicates the previously used cut-off (Prigozhin and Krasileva 2021) to determine highly variable residues (Entropy = 1.5 or normalized entropy = 0.347). **D.** The distribution of the normalized Shannon entropy across the leucine-rich repeat (LRR) domain of the Sr50 group. The LRR region of the normalized Shannon entropy for the Sr50 group provided in C is depicted as a heatmap to provide a closer view of sequence variations. Each LRR unit and each position within the LRR unit was defined according to the general motif LXX**LXL**XX(C/N).

To incorporate the location of highly variable sites within the NLR structure, we created a plot displaying Shannon entropy measures of the residues within each leucine-rich repeat unit that form the beta sheets that form the inner concave surface of the LRR. We noticed that LRRs 8-14 contained a cluster of sites with high Shannon entropy that we hypothesized to be a potential effector binding pocket (Fig. 1D). Since high entropy plots of the four phylogenetic groups were remarkably similar, we created the same plot displaying high Shannon entropy measures in key residues of the LRR units (Fig. S1). High Shannon entropy scores occur similarly towards the end of the LRRs in units 8-16 in all four phylogenetic groups. This suggests more specifically. that the effector binding pockets are in the LRR same region across homologous NLRs, despite their ability to recognize structurally distinct effector. These results indicate that highly variable sites within the NLRs of the MLA family could be where diversification of paralogs occurs.

To test our prediction, we created a chimeric protein consisting of Sr33 and the LRRs 8-14 of Sr50 that we will refer to as Sr33/50 throughout. A schematic illustrating the location of the transferred region within the chimera can be found in Figure S2. To test whether the chimera Sr33/50 is capable of recognizing AvrSr50 and subsequently inducing cell death, we utilized a wheat protoplast cell death assay, as previously described (Saur et al., 2019; Yoshida et al., 2009). In this assay, wheat protoplasts are transfected with luciferase and a combination of NLRs and effectors. A positive interaction between NLR and effector leads to cell death as exemplified by a lack of luciferase activity 2 days after transfection. Co-transfection of Sr50 with AvrSr50 led to cell death and strongly reduced luminescence, whereas co-transfection of Sr33 with AvrSr50 did not lead to a reduction as compared to NLRs alone (Fig. 2A). The chimeric protein Sr33/50 was able to recognize AvrSr50 as demonstrated by the strong reduction of luminescence signifying cell death (Fig. 2A). This result confirmed the prediction that LRRs 8-14 mediate effector recognition in Sr50 because the transfer of these LRRs to Sr33 was sufficient to transfer effector specificity.

**Fig 2:**
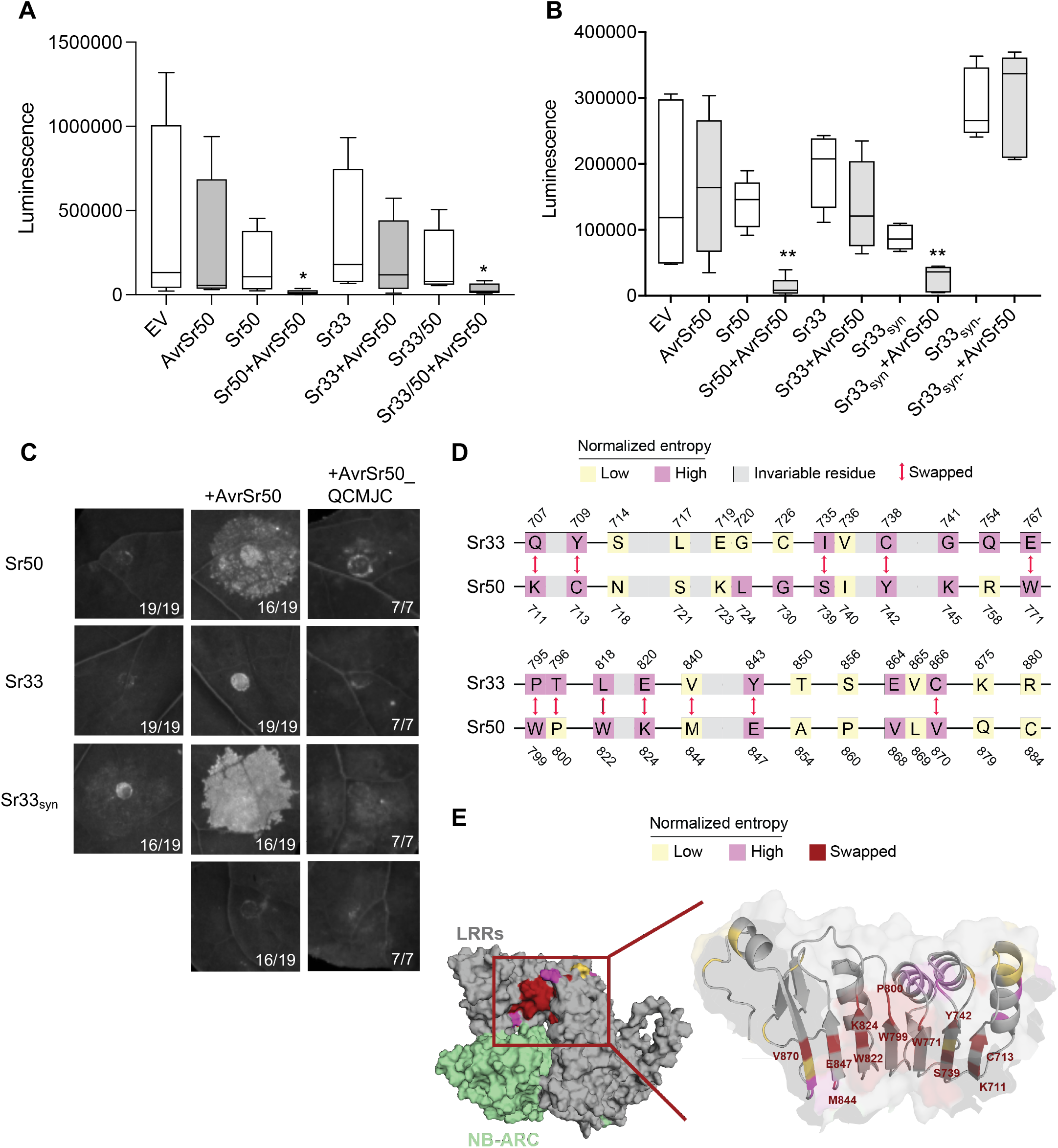
Transfer of domains and amino acid groups from Sr50 to Sr33 enables the recognition of the previously unrecognized effector AvrSr50. **A.** The chimeric protein Sr33/50 recognizes AvrSr50 and induces cell death in the wheat protoplast cell death assay. Graph shows data from four independent biological replicates and asterisk indicates significant difference between EV and AvrSr50 treatment (paired t-test, n=4, *p < 0.05). Seedlings of the wheat cultivar Fielder were peeled and digested in enzyme solution for protoplast generation. After successful protoplast isolation they were transfected with constructs for the wheat protoplast assay as described previously (Saur et al., 2019). Protoplasts were harvested 2 days later and luminescence was measured in a TECAN Infinite F Plex plate reader utilizing the Promega Luciferase Assay System (Promega, Cat. #E4550). **B**. Twelve amino acid changes on Sr33 are sufficient to enable recognition of AvrSr50 as exemplified by the synthetic NLR Sr33_syn_. We introduced twelve amino acid changes into the NLR Sr33 that is unable to induce cell death in wheat protoplasts in response to AvrSr50 and named the new NLR Sr33_syn_. In contrast to Sr33, Sr33_syn_ is able to recognize and execute cell death in an AvrSr50-dependent manner. The synthetic NLR Sr33_syn-_ that only contains 11 amino acid substitutions does not recognize AvrSr50. Graph shows data from three independent biological replicates and asterisk indicates significant difference between EV and AvrSr50 treatment (paired t-test, n=3, **p << 0.01) **C**. Sr33_syn_ induces cell death in response to AvrSr50 in *N. benthamiana*. None of the NLRs initiated cell death in response to the effector variant AvrSr50_QCMJC. Leaves of 4-5 weeks old *N. benthamiana* plants were infiltrated with *A. tumefaciens* carrying NLRs and effectors at OD_600_=0.3 and 0.4, respectively. UV images were taken on a BioRad Gel Imager 3-5 days after infiltration. Numbers on images correspond to the numbers of leaves that showed the depicted phenotype. **D**. Amino acid residues in Sr33 and Sr50 targeted for swapping. The amino acid sequences and positions in LRRs 8-13 are depicted if the sequences are characterized by substitutions between Sr33 and Sr50. Low and high entropy were determined based on the cut-off (0.347) and are indicated in different colors. The 12 amino acid sequences used to generate Sr33_syn_ are indicated as ‘swapped’. **E**. Variable amino acids mapped to the Sr50 structure. The structure of Sr50 was predicted with AlphaFold (Jumper et al. 2021), and only the NB-ARC domain and LRRs are shown. The residues targeted for swapping are annotated.

We then aimed to identify the individual amino acids involved in the recognition of AvrSr50. In total, there are 26 amino acid polymorphisms in the LRRs 8-14 between Sr50 and Sr33. We mapped the variable residues onto the surface of Sr50 and Sr33 and selected 13 amino acids that fulfilled most of these criteria: high Shannon entropy signature, differing amino acid side chain property between Sr33 and Sr50 or exposure at the surface of the beta-sheets that face the NB-ARC domain (Fig. 2D and E). We synthesized several iterations of Sr33 carrying mutations in these predicted amino acid residues (with 10 to 13 amino acid substitutions) that correspond to the hypothesized binding sites in LRRs 8-13 of Sr50 and tested these synthetic NLRs in wheat protoplasts. Among the constructs we tested, the synthetic NLR Sr33_syn_ carries 12 selected amino acid polymorphisms (Fig. 2D) and initiated cell death in wheat protoplasts in an AvrSr50-dependent manner while Sr33_syn-_ did not (Fig. 2B). These two synthetic NLRs differ only in one amino acid residue that seems to affect recognition. While Sr33_syn_ carries a methionine in LRR13 (M840), the NLR Sr33_syn-_ carries a valine residue in the respective position (V840) (Fig. 2D, Fig. S3D). Our observation is therefore that 12 amino acid residues were the minimal amount needed to transfer recognition of AvrSr50 to Sr33. To substantiate this observation, we infiltrated these mutants in *N. benthamiana* with AvrSr50 to test if these mutations enable Sr33 to recognize AvrSr50 and induce cell death. Our results demonstrate that the transfer of 12 amino acids is sufficient to alter recognition specificity of Sr33 as it shows a clear cell death response when co-infiltrated with AvrSr50 (Fig. 2C). In contrast, the effector variant AvrSr50_QCMJC that evades recognition by Sr50 is also not recognized by Sr33syn, demonstrating that the gained effector recognition is AvrSr50-specific. These results demonstrate that we successfully predicted amino acids involved in the recognition of AvrSr50 through Sr50 and that effector recognition can be transferred to a closely related NLR.

In our protoplast cell death assays, we noticed that Sr50 could exhibit autoactivity whereas Sr33 and Sr33/50 did not. Furthermore, it has been published that Sr50 and Sr33 can exhibit autoactivity when over-expressed in *N. benthamiana*, but not when expressed under the weaker, native Sr50 promoter (Cesari et al., 2016). We observed the same in *N. benthamiana* where expression with *p35S* caused autoactivity whereas expression driven by a previously described, weaker *pRPP13* (Rentel et al., 2008) greatly reduced cell death in the absence of effector (Fig. S3). However, in wheat protoplasts only Sr50 exhibited varying levels of autoactivity while Sr33 consistently did not (Fig. 3A). Since the two NLRs are highly similar, we reasoned that we could reduce the Sr50 autoactive phenotype by changing certain amino acids to the corresponding sites in Sr33. In the iterations of synthetic Sr33 NLRs we observed that many of them exhibited autoactivity. Together with our data from Sr50, Sr33 and Sr33_syn_ we selected two residues to be mutated in Sr50 that we hypothesized to affect the autoactivity phenotype in wheat protoplasts. We tested the autoactivity of the Sr50 mutants in the wheat protoplast cell death assay (Fig. 3A) and observed reduced autoactivity of NLR alone while preserving effector recognition (Fig. 3A). We also tested whether effector recognition is retained and remains specific in *N. benthamiana*. We infiltrated Sr50 and the single and double mutants with both AvrSr50 and AvrSr50_QCMJC. All Sr50 variants induce cell death in an AvrSr50-dependent manner, while no cell death was observed with the AvrSr50_QCMJC effector variant (Fig. 3B). We therefore show that closely related NLRs can also provide information about activation mechanisms and be utilized to reduce autoactivity without affecting effector recognition specificity

**Fig 3:**
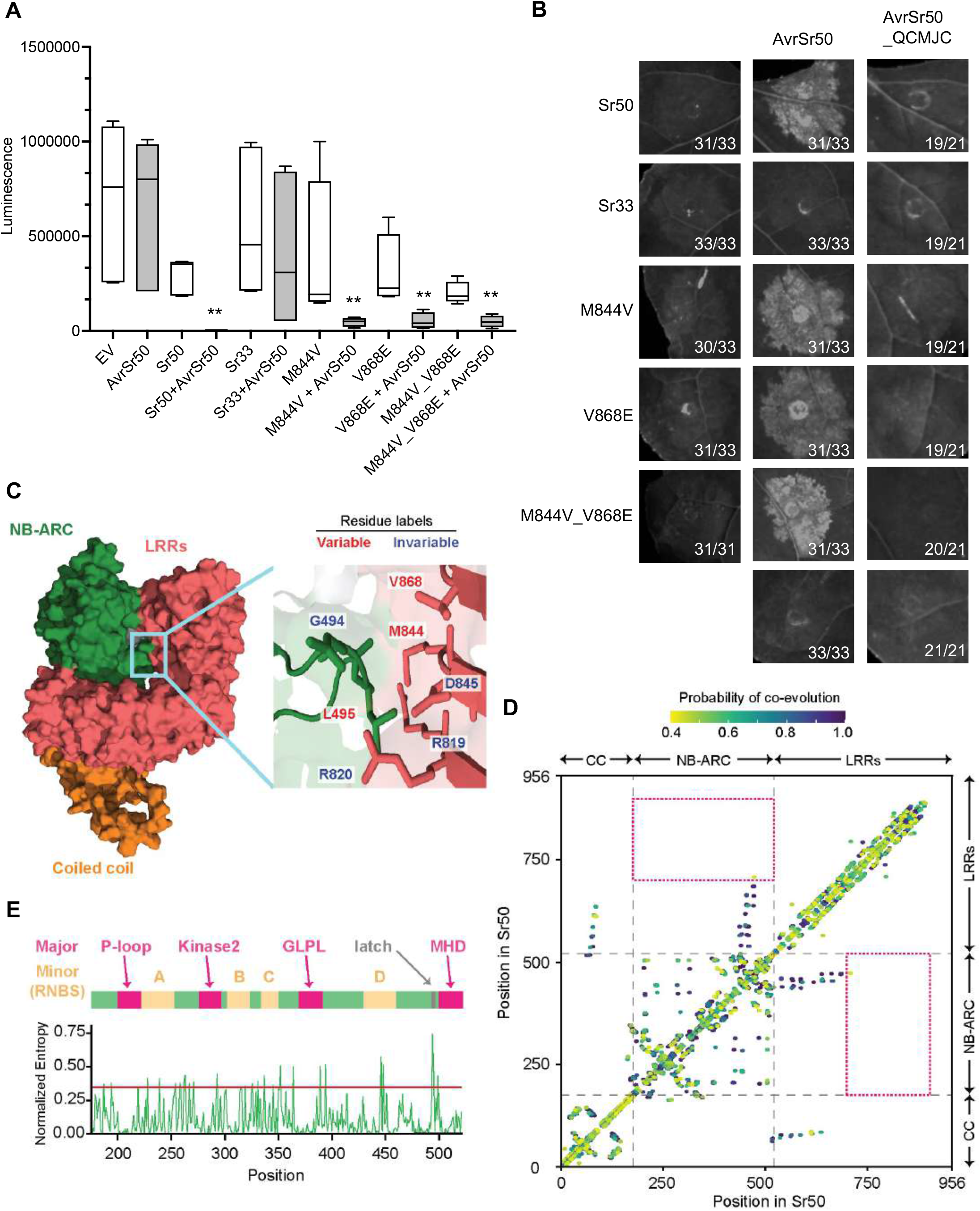
Sr50 autoactivity is decreased through mutations in its LRR. **A**. Single and double mutants of Sr50 show decreased autoactivity in wheat protoplasts compared to wild type Sr50. Graph shows data from three independent biological replicates and asterisk indicates significant difference between EV and AvrSr50 treatment (paired t-test, n=3, **p << 0.01). We introduced single and double amino acid changes into the NLR Sr50 to decrease its autoactivity. The sites changed were changed to the corresponding amino acids in that position within Sr33 as this NLR does not exhibit the same autoactivity phenotype in wheat protoplasts. **B**. Single and double mutants of Sr50 induce cell death in *N. benthamiana* in response to AvrSr50. None of the NLRs initiated cell death in response to the effector variant AvrSr50_QCMJC. Leaves of 4-5 weeks old *N. benthamiana* plants were infiltrated with *A. tumefaciens* carrying NLRs and effectors at OD_600_=0.3 and 0.4, respectively. UV images were taken on a BioRad Gel Imager 3-5 days after infiltration. Numbers on images correspond to the numbers of leaves that showed the depicted phenotype. **C**. The Sr50 structure and insert to the NB-ARC latch. Structure was generated using AlphaFold. NB-ARC, LRR and CC domains are shown in green, red and orange, respectively. Blue box enlarges contact point between NB-ARC and LRR domains and the involved amino acids; we refer to this interaction as NB-ARC LRR latch. Some of the residues in this NB-ARC LRR latch are depicted. If a homologous position of the residue is substituted in Sr33, the residue is indicated as ‘variable’. Otherwise, it is indicated as ‘invariable’ **D**. The signature of inter-residual co-evolution captured by Gremlin. The filtered multiple sequence alignment (MSA) collected during structure prediction with AlphaFold for Sr50 was used to infer inter-residual co-evolution with Gremlin. The data provided in Table S5 is visualized. Only the pairs with probability ≥ 0.40 were indicated. If inter-residual co-evolution was detected, a colored dot is indicated in the corresponding position of the pair. The x- and y-axis indicate amino acid position in Sr50. Multi-domain architecture (Coiled coil (CC), NB-ARC, and leucine-rich repeats (LRRs)) is provided around and across the MLA. The red dashed boxes indicate where coevolution between the NB-ARC domain and terminal LRRs would appear. **E**. The sequence feature of Sr50’s NB-ARC domain. The normalized entropy was calculated with all MLA members used in the study. Major and minor motifs within the NB-ARC are shown in relation to normalized entropy values. Gray arrow indicates amino acid involved in NB-ARC/LRR latch interaction.

To understand why certain amino acid sites had such a drastic effect on Sr50’s autoactivity, we modeled the structure of Sr50 with AlphaFold. The model showed that LRR residues M844 and V868 are in close contact with the NB-ARC domain, and we hypothesize that these residues are important in maintaining the inactive state of the NLR through the interaction with the NB-ARC domain (Fig. 3C). Upon calculation of the pairwise residual distances in the predicted structure, we found that two residues in the NB-ARC domain were in proximity with terminal LRR residues: G223 and L495 (Fig. S6). Among the two residues in the NB-ARC, only L495, that appears in a short stretch of random coils, was in contact with LRRs 8-13 (< 7.5 Å) (Fig. 3C; Fig. S6). We named the residues that define this particular inter-domain interaction the NB-ARC/LRR latch (Fig. 3C). Interestingly, of the specified seven residues involved in the NB-ARC/LRR latch only three were polymorphic between Sr50 and Sr33 (indicated as variable in Fig. 3C) and we identified two of these in our functional study of Sr50 mutants. Moreover, the residue M844 was required for effector recognition in Sr33syn, as the variant Sr33_syn-_ lacks this methionine residue and did not recognize AvrSr50 (Fig. 2B, 2C; Fig. 2D). These results suggest that LRR residues within Sr50 and Sr33_syn_ that mediate effector binding also affect receptor autoactivation.

Since the change of a single amino acid resulted in such a large change of the autoactive phenotype, we were curious if there might be any co-evolutionary signatures between residues in the NB-ARC/LRR latch. Co-evolutionary inference relies on a deep MSA composed of diverse homologs. We therefore constructed an MSA with 21,518 putative CNLs from 330 Viridiplantae species and tested co-evolution with direct coupling analysis (DCA) (Zerihun et al., 2020) and Gremlin (Ovchinnikov et al., 2014). When we mapped the inference to the Sr50 sequence we found that residues located close to each other generally displayed co-evolutionary signature (Fig. 3D; Table S5). However, the NB-ARC domain and LRRs 8-13 (red dotted areas in Fig. 3D) lacked signal of coevolution, although the residues in the NB-ARC/LRR latch are physically close (< 7.5 Å, Fig S6). We also investigated whether the sites in the NB-ARC that are in proximity to the LRR latch correspond to any known, highly conserved major or minor motifs in the NB-ARC domain and found that they did not (Fig. 3E). For the MLA family member, although sequence conservation of the NB-ARC latch was not as high as that of the major motifs, glycine residue (G496) was highly conserved (Fig. 3C; Fig. 3E). We postulate that this glycine is important to make a sharp turn in the protein’s tertiary structure and give rise to the structural characteristic of the NB-ARC latch. Together, our functional and structural analyses highlight a single LRR region, where residues which are in close physical proximity to the NB-ARC have the dual function in constraining autoactivity and enabling effector specificity.

## Discussion

A large number of resistance genes deployed are NLRs, which makes them an excellent target for improvement of plant health. A bottleneck in the rational design of new NLR alleles has been the lack of understanding of effector binding and subsequent receptor activation. Though it has been known for a long time that the LRR domains play a crucial role in effector specificity (Ellis et al., 1999; Shen et al., 2003; Krasileva et al., 2010), difficulties in structural biology have limited our ability to close in on the specific sites mediating effector binding. To successfully generate new effector recognition specificities, we need to decipher sites within NLRs that mediate effector binding and those required to maintain stability and the inactive state. Recent advances both in structural biology (Martin et al., 2020; Wang et al., 2019a, 2019b) and NLR engineering (Cesari et al., 2021; De la Concepcion et al., 2019; Pottinger et al., 2020) have exemplified success in the engineering of NLR proteins.

With the increasing amount of genomic data of many different species and populations to analyze NLR evolution, it became possible to predict and engineer effector binding specificity. It was previously shown that the sequence diversity data and Shannon entropy can be used to predict if an NLR is a direct or indirect binder and where within the NLR effector binding occurs (Prigozhin and Krasileva, 2021). Here, we combined Shannon entropy with structural modeling to predict a potential effector binding site and guide rational engineering to transfer effector recognition specificity between distantly related homologs. In our structural analyses, we initially utilized PHYRE2 (Kelley et al., 2015) to model the Sr50 structure but the homologous template-based model was incomplete as the template structure is not fully resolved and the resulting model was therefore missing several leucine-rich repeats. Since it was the only model available at the time, we used the incomplete model of the LRR repeats to the extent possible to identify surface-exposed sites for potential engineering. After the release of AlphaFold (Evans et al., 2021; Jumper et al., 2021), we recreated the Sr50 protein model and found that the LRRs of both models were consistent. By mapping Shannon entropy values and amino acid polymorphisms between Sr50 and Sr33 on the exposed beta-sheets of the LRR domain, we were able to use additional criteria to select residues for engineering as opposed to using just Shannon entropy scores alone. This approach greatly enhanced our ability to successfully engineer Sr33 as exemplified by the identification of the residue V840. Its corresponding site M844 in Sr50 has a low Shannon entropy score but was required for the transfer of effector recognition in Sr33_syn_. We therefore show that the application of several analyses can be complementary and help successfully engineer NLR recognition specificities even in the absence of experimentally determined structural data.

The ‘on’ and ‘off’ states of NLRs are tightly controlled and recent cryo-electron microscopy structures of an indirect binder NLR, *Arabidopsis thaliana* HOPZ-ACTIVATED RESISTANCE 1 (ZAR1), confirmed what has previously been inferred through domain interaction and swap data: the inactive and ADP-bound state was monomeric and had multiple intramolecular contact points between the LRR and NB-ARC (Wang et al., 2019a, 2019b). Moreover, previous studies showed that the correct interactions of NB-ARC and LRR are required to avoid autoactivity of the NLR (Qi et al., 2012; Rairdan and Moffett, 2006; Slootweg et al., 2013) and even effector responsiveness (Ma et al., 2018). The differing autoactivity phenotypes of Sr33, Sr50 and our Sr33_syn_ variants allowed us to investigate this as the differences among all three proteins was contained to their LRR domains. The data from this study supports the notion that interactions between NB-ARC and LRR are crucial in maintaining the inactive state of the direct binder NLR Sr50. A recent paper suggested that the interaction between the NB-ARC and LRR domains serves the purpose of preventing NLR oligomerization. (Zhao et al., 2021). It would be interesting to see if residues in the NB-ARC/LRR latch also play a role in NLR oligomerization or whether the interaction is merely to prevent access to the sites mediating nucleotide exchange and oligomerization.

Through our introduction of amino acid changes in Sr50 and structural modeling, we identified a direct interaction point of NB-ARC and LRR that we termed NB-ARC/LRR latch. This interaction most likely results in a conformation of the NLR that hinders the exchange of ADP to ATP in the NB-ARC domain or oligomerization into resistosomes. Strikingly, our data suggests that control of autoactivity and effector specificity are located within the same region of the NLR and mediated by the same residues. We hypothesize that the effector competes with and disrupts the interaction of the NB-ARC/LRR latch leading to the activation of the NLR. Our data suggests that this interaction can also be disrupted through the exchange of amino acid residues in the NB-ARC/LRR latch. We therefore hypothesize that the disruption of NB-ARC/LRR latch is a crucial step in NLR activation and can be caused by effector binding or unstable intramolecular interactions. We therefore show that the strength of the NB-ARC/LRR latch interaction is an important criterion to consider when undertaking NLR engineering approaches.

It has been proposed that NLRs exist in fluent states of active and inactive and effector binding stabilizes the active state thereby shifting the equilibrium in its favor – this is referred to as the equilibrium-based switch model (Bernoux et al., 2016). We propose that the activation of hvNLRs, such as Sr50, follows the equilibrium-based switch model (Bernoux et al., 2016) based on our observations of expression-dependent autoactivity and our ability to change this phenotype with a few amino acid substitutions. According to this model there is an equal pool of activated and inactive receptors in each cell in the absence of a trigger. Effector binding stabilizes the activated conformation and leads to the activation of downstream signaling pathways. This implies that the cell can cope with a certain amount of activated receptors and overexpression would lead to an excess of activated receptors. This could push the resulting cellular changes over the threshold required to initiate the cell death response, for instance through increased calcium influx caused by the insertion of CC domains of activated, oligomerized receptors into the membrane. Collectively, our data, which showed that Sr50 exhibits varying degrees of autoactivity depending on its expression level (Fig. 3A, Fig. S3), suggests that this NLR follows the equilibrium switch model of activation. It will therefore be crucial to understand the limitations of deploying NLRs that follow this model of activation and ensure the use of appropriate promoters in the generation of new resistant plants with hvNLRs. We hypothesize that having multiple factors that can drastically change autoactivity phenotypes such as expression levels and intramolecular interaction strengths is what allows for the rapid generation of diversity in hvNLRs.

It is tempting to hypothesize that the amino acid sites we identified in the latch would evolve together to maintain the rate at which the NLR can be activated. However, neither of the analyses we performed detected any co-evolutionary signature and even suggested that co-evolution between NB-ARC and terminal LRRs is lacking. Since there are at least seven residues involved in the NB-ARC/LRR latch, it is conceivable that any co-evolutionary signature would be too diluted among all involved residues to be detected or that the activation mechanism of the NLR is too complicated to be summarized by just three residues. The fluidity of Sr50’s activation state is another possible reason for the lack of measurable co-evolution, as the ‘on’ and ‘off’ states of this NLR are affected by many different factors. Moreover, the co-evolutionary analyses that are currently at our disposal require rather large datasets with an evolutionarily broad range of NLRs. If the co-evolution of the NB-ARC/LRR latch is lineage-specific, and multiple modes of interactions exist for these inter-domains, the datasets may obscure any co-evolutionary signatures. Elucidating the co-evolution of the NB-ARC/LRR latch will require better understanding of how NLRs are cycled between active and inactive states (there is no experimental structure of an inactive direct binder NLR), how many different modes there may be, and which NLRs follow each mode of activation.

This study shows a major advance towards the successful engineering of new effector recognition specificities in direct binder NLRs. We demonstrate that effector binding sites predicted through diversity analysis can be successfully used to engineer or transfer recognition specificities. Our study also demonstrates that for the successful engineering of an immune receptor we also need to identify sites that are important for maintaining the inactive state of the NLR. In addition to the generation of novel NLRs, we show that existing NLRs can be engineered to recognize a new effector and more importantly recognition can be transferred between closely related NLRs. This means we might be able to generate resistant plants through precise genome editing of NLR homologs. Through this approach we can use existing NLRs that are available for deployment through map-based cloning approaches and edit their homologs in elite crop germplasms. The region required for the transfer of effector recognition involved as little as 12 amino acid substitutions within 450 bp. When this region were to be introduced into a closely related homolog through homologous recombination the engineered plants would follow the regulatory exemptions described in the SECURE rule from the USDA, given this variation is already present in the edited plant’s gene pool (Hoffman, 2021). The ability to transfer effector recognition between close homologs also gives us the ability to avoid interspecies transfer of genes, which is controversial for some opponents of transgenic plants. The information provided in this study may therefore help to drive forward the generation of resistant elite crop germplasm.

## Materials and Methods

### Plant materials and growth conditions

*Nicotiana benthamiana* plants were grown in a Conviron growth chamber with 16 hours of 150 /s light at 24°C. *Triticum aestivum* cultivar Fielder seeds were sterilized in 10% bleach solution for 20 minutes. They were placed on water and filter paper in a plastic cup and the cup was sealed by placing another cup on top and adhering them together with Millipore tape. The cups were kept on the west-facing windowsill for 7-14 days, depending on the season.

### Cloning and vectors

All vectors cloned for this study were cloned following the Golden Gate cloning technique (Weber et al., 2011; Werner et al., 2012) by utilizing the vectors deposited as MoClo Plant Parts (#1000000047) and Tool kits (#1000000044) at Addgene. For expression in wheat protoplasts, all genes were driven by the *pZmUBQ* promoter (pICSL12009) and the *tNos* terminator (pICH41421), whereas for expression in *N. benthamiana* the *pRPP13* promoter (Rentel et al., 2008) and *tNos* terminator, as well as *p35S* (pICH51277) and *t35S* (pICH41414) were used. The following genes were domesticated for Golden Gate by removing internal restriction sites for *BsaI* and *BpiI* and were cloned into the corresponding level 0 acceptor: Sr33, Sr50, AvrSr50, AvrSr50_QCMJC, and pRPP13. All coding sequences were cloned both with and without stop codon. Mutants of Sr33 were created by synthesizing the LRRs 8-13 with 10 amino acid substitutions and ligating it together with PCR fragments of the start and end of Sr33. Additional mutations were added using site-directed mutagenesis PCR, by designing two overlapping primers with the desired mutation in the middle, amplifying the vector by Phusion PCR and digesting the template with *DpnI*. Mutants of Sr50 were created by site-directed mutagenesis PCR. All vectors and primers used in this study are described in Table S5 and S6.

### Wheat protoplast isolation, transfection, and cell death assay

Wheat protoplast isolation and transfection was adapted from (Saur et al., 2019; Yoshida et al., 2009). The epidermis of 7-14 days old seedlings was peeled off by making a shallow cut in the adaxial epidermis with a sharp razor blade, folding the leaf over at the cutting edge and pulling back the leaf tip with the abaxial epidermis. The peeled leaf sections were placed next to each other on masking tape with the abaxial side facing upwards and placed into enzyme solution facing down. Washing and transfection were performed as previously described. Protoplast number was counted with a hemocytometer and adjusted to 300,000 protoplasts/mL. All plasmids were transfected at a concentration of 15 μg and an empty vector was added to ensure all protoplasts were transfected with the same final DNA concentration. After transfection, protoplasts were incubated in a BSA-coated 12-well plate at room temperature in the dark for 48 hours. Subsequently, protoplasts were collected by centrifugation at 100g for 3 minutes. Supernatant was discarded and protoplasts resuspended in 2x Cell Culture Lysis Reagent (Promega, Cat. #E1531) and incubated on ice for 20 minutes in the dark. The lysed cell contents were centrifuged at 1,000g for 3 minutes and divided into a white 96-well plate (Greiner Bio-One, Cat. # 655075) in triplicate. An equal amount of Luciferase Assay Substrate (Promega, Cat. #E4550) was added and luminescence was measured in a TECAN Infinite F Plex plate reader (20 minutes, settle time 100 ms, no attenuation).

### Agrobacterium-mediated transformation of *N. benthamiana*

*Agrobacterium tumefaciens* GV3101:pMP90 transformed with the corresponding construct were grown in LB medium (supplemented with 50 μg/ml Rifampicin, 25 μg/ml Gentamycin and 100 μg/ml Carbenicillin) at 28°C. The cultures were centrifuged at 3,000 g for 15 minutes and the pellet resuspended in infiltration medium (10 mM MES pH=5.6, 10 mM MgCl2). Optical density was adjusted to a final OD_600_=0.3. If several constructs were co-infiltrated the suspensions of individual constructs were mixed in a 1:1 ratio while ensuring that all constructs reached a final density of 0.3. Leaves of 4-5 weeks old *N. benthamiana* plants were inoculated with the suspension using a blunt syringe. Pictures of leaves were taken 2-3 days after inoculation.

### Manual curation of NLRs

To ensure sequence diversity needed to calculate entropy, we manually curated a subset of NLRs from 14 wheat genomes (Walkowiak et al., 2020) (Table S1). From the wheat reference genome, we first identified the NLRs that belong to the MLA family, guided by the previous study (Steuernagel et al., 2020). Their NB-ARC domain was searched with TBLASTN (-evalue 1E-10) (Camacho et al., 2009) against the 14 wheat genomes. We identified matches that showed 85% or more sequence identity against the queries and extracted the putative NLR loci with 10,000 flanking regions on both ends. We used MAKER v3.01.04 (Cantarel et al., 2008) to obtain gene models in these regions, relying on two ab initio predictors, Augustus v3.4.0 (--species wheat) (Stanke et al., 2006) and GeneMark v4.68_lic (Brůna et al., 2020), as well as CDS and protein sequences of the wheat reference genome annotation. The gene models were loaded into Apollo genome browser v2.0.6 (Dunn et al., 2019) and manually curated. The manual curation generated many redundant NLRs from the 14 wheat genomes, the sequences of which are nearly identical. The redundant sequences can inflate their weighting and copy numbers in entropy calculations and the phylogenetic analyses, which can mislead evolutionary conclusions. We therefore reduced redundancy of the wheat NLRs based on their sequence identity (95%) with CD-hit (-c 0.95) v4.8.1(Li and Godzik, 2006).

### Phylogenetic analyses

To infer a species tree of the selected Pooideae members, we first identified complete BUSCO genes present in all nine species with BUSCO v5.2.2 (Seppey et al., 2019) and the poales_odb10 database and randomly selected 500 (Table S1). Each set of orthologous protein sequences was aligned with MAFFT v7.487 (--maxiterate 1000 --globalpair) (Katoh and Standley, 2013). All MSAs were concatenated, trimmed with TrimAl v1.4.rev15 (-gt 0.3) (Capella-Gutiérrez et al., 2009) and used to infer a phylogenetic tree with FastTree v2.1.10 (Price et al., 2010).

To infer the MLA family tree, we collected the MLA family members from the Pooideae species, filtered wheat NLR annotation sets and experimentally validated NLRs (Kourelis et al., 2021), guided by the classification of the previous study (Steuernagel et al., 2020). Their full-length sequences were aligned with MAFFT, and we trimmed out all columns where both Sr33 and Sr50 contained gaps. The resulting MSA was used to infer a phylogenetic tree with FastTree (-slow). Sr35 was used as an outgroup to root the tree.

### Protein structure prediction and structural analysis

We used AlphaFold v2.0.0 to predict the structure of Sr50 (Jumper et al., 2021; Evans et al., 2021). The homologs were collected from the full databases, and homologous templates downloadable by July 20th, 2020, were included in the template database. Models 1, 3, 4 and 5 as well as ptm model 2 were used for structural inference. The best model was selected as a final structure based on the average predicted lDDT scores. We calculated the distances between residues in the predicted structure with Biopython (Cock et al., 2009). The structure was visualized through PyMOL v2.5.

### Co-evolutionary analyses

To collect CNLs for co-evolutionary analyses, we extracted the protein sequences from the Viridiplantae species from the uniref100 database (2022-0222). The gene models containing the NB-ARC domain (PF00931) were identified as putative NLRs with hmmsearch 3.1b2 (--domE 1e-4 -E 1e-4) and annotated with InterProScan v5.52-86.0 (--appl Pfam-33.1) (Eddy, 2011; Jones et al., 2014). Any sequences with other PFAM entries than NB-ARC domain, Rx N-terminal domain (PF18052) and Leucine Rich Repeat (PF00560, PF07725 and PF13855) were excluded. The NB-ARC domain of the remaining NLRs were aligned with MAFFT (--auto --global), trimmed with TrimAl (-gt 0.3) and used to infer a phylogenetic tree with FastTree. We identified remaining TNL and RNL clades and removed the clade members.

We searched this database with Sr50 as a query with jackhmmer (--domE 1e-10 -E 1e-10 -N 5). The MSA was trimmed with respect to Sr50, and homologs without ≥ 75% coverage were removed. We used this filtered MSA for DCA (Zerihun et al., 2020) and Gremlin (Ovchinnikov et al., 2014). The pseudolikelihood maximization algorithm of the pydca package was run for DCA (--max_iterations 1000 --apc). The filtered MSA was also submitted to the online Gremlin server (HHblits iterations = 0, Coverage = 50, Remove gaps = 50).

### Shannon entropy calculation

We computed normalized Shannon entropy 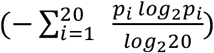 where *p_i_* is the probability of observing an amino acid, i, out of the twenty in the given position of the input MSA. Gaps were ignored in entropy calculation. However, if the position of the MSA was composed of 50% or more gap characters, the entropy was not calculated. The normalized Shannon entropy ranges from 0 (no variability) to 1 (maximum variability). The normalized entropy score of 0.347 was used to define highly variable residues, following the previous study (Jumper et al., 2021; Prigozhin and Krasileva, 2021).

## Data Availability

All data and scripts used in this study are available in Github: https://github.com/krasileva-group/Sr33-Sr50_analysis.

## Acknowledgements

We thank Dr. Isabel Saur and Salome Wilson for helpful practical advice on the protoplast cell death assay. We thank Dr. Daniil Prigozhin for initial Shannon entropy analyses and effector binding site prediction on Sr50, and helpful comments on the manuscript. JAT is grateful for many personal discussions with Dr. Erin Baggs about this project and thoughtful suggestions on the bioinformatics analyses. We are grateful to all members of the Krasileva lab for thoughtful comments and discussions of the presented material. The project has been funded by Gordon and Betty Moore Foundation, the Foundation for Food and Agriculture and 2Blades Foundation. KS has been supported by the Berkeley Graduate Fellowship.

## Supplement

**Figure S1.**
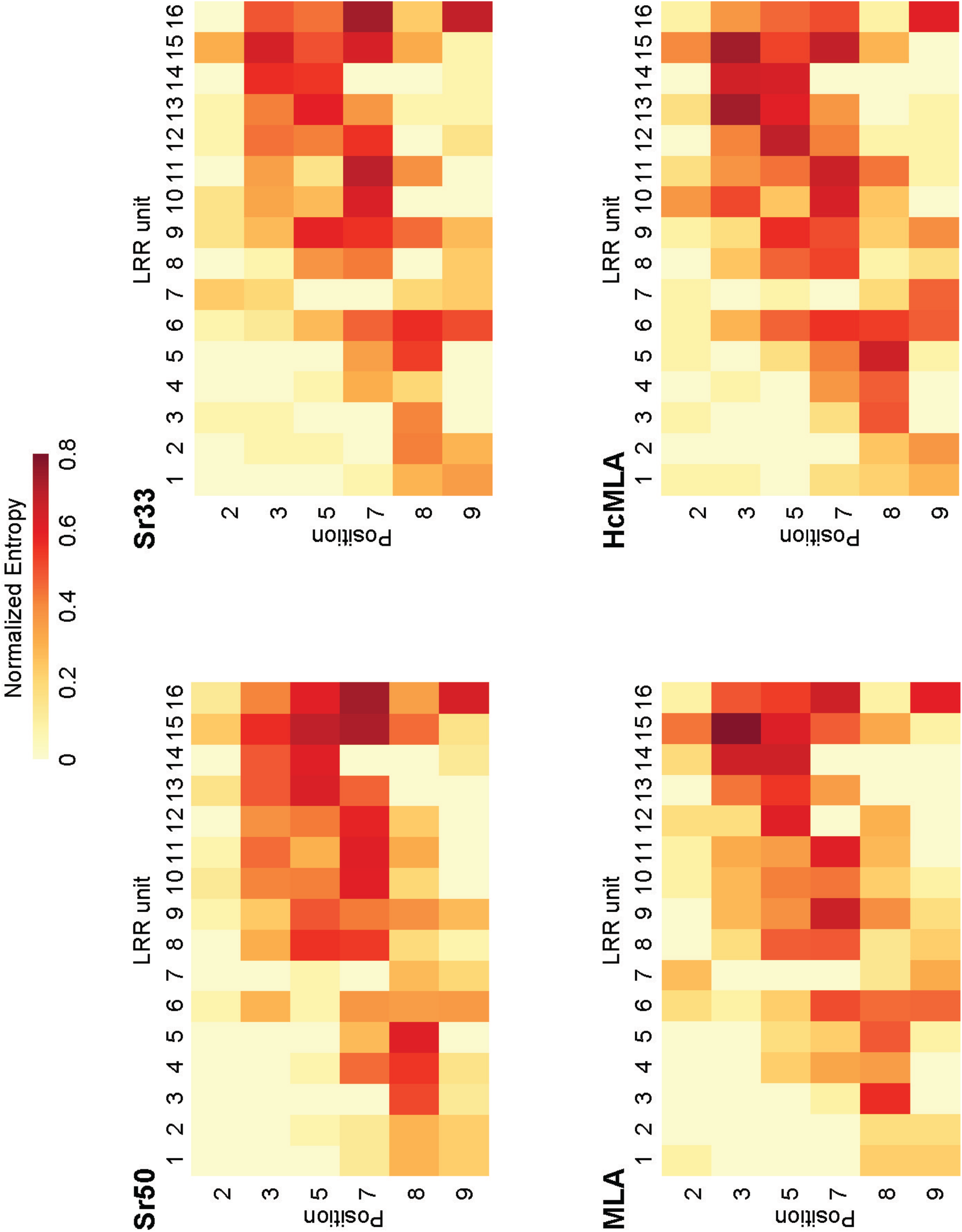
The distribution of the normalized Shannon entropy across the leucine-rich repeat (LRR) domain of the four phylogenetic groups from Fig. 1A. The normalized Shannon entropy across LRRs 1-16 for each phylogenetic group is depicted as a heatmap to provide a closer view of sequence variations. Each LRR unit and each position within the LRR unit was defined according to the general motif LXX**LXL**XX(C/N).

**Figure S2.**
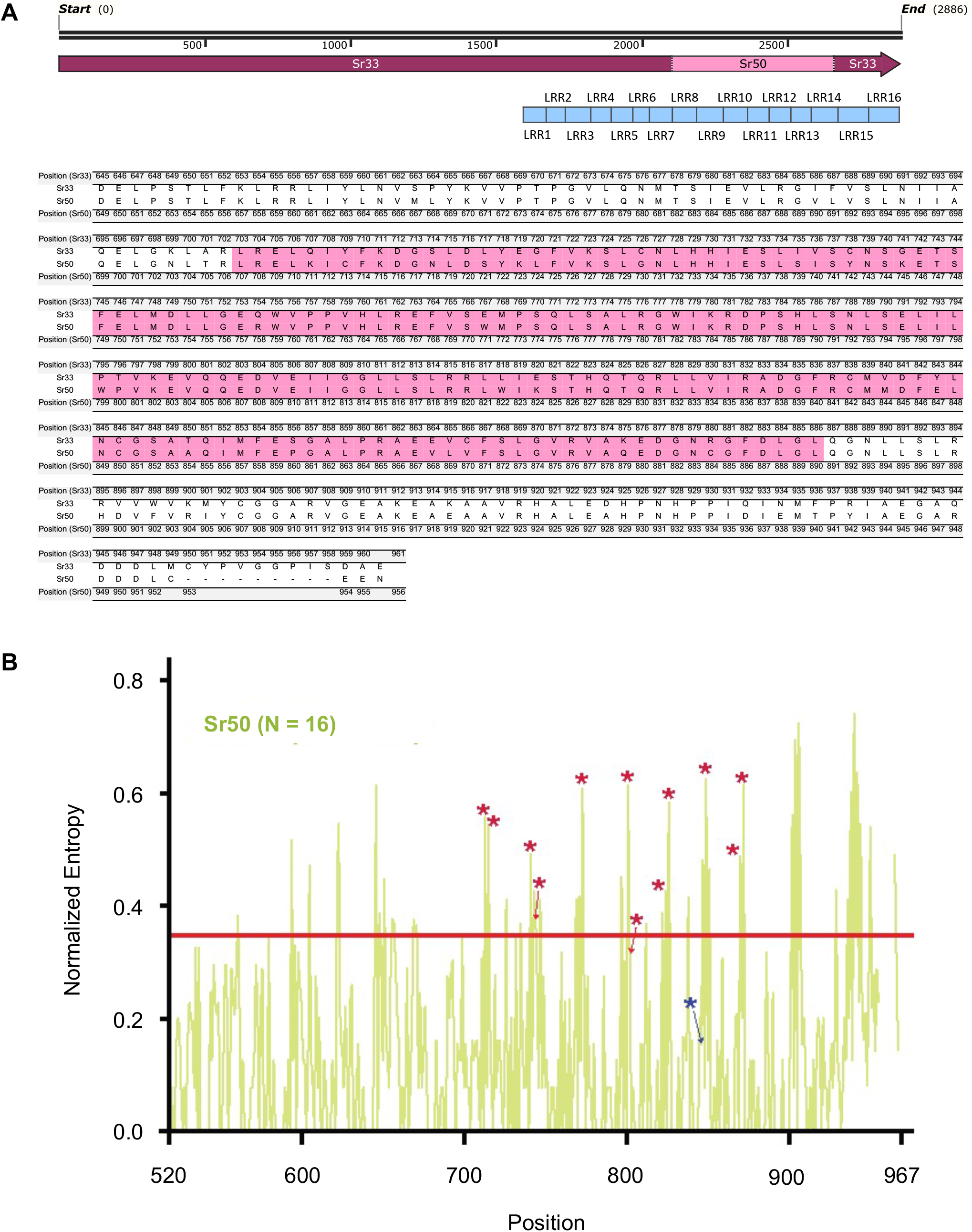
Schematic representation of swapped the region in the chimeric NLR Sr33/50. **A.** The chimeric protein Sr33/50 was created by replacing amino acids in the region of LRRs 8-14. The NLR consists of Sr33 ranging from AA 1-702 and 887-961. In place the region 703-886 are the corresponding sites in Sr50. **B**. Normalized Shannon entropy of NLRs in the Sr50 clade. The asterisks indicate amino acids that were utilized in the generation of Sr33_syn_. Red asterisks indicate positions in which either Sr33 or Sr50 has show high entropy peaks determined by the cut-off of 0.347.; The blue asterisk is the position in which both Sr33 and Sr50 had low entropy values and corresponds to the low entropy peak (M844) that governs autoactivity within Sr50.

**Figure S3.**
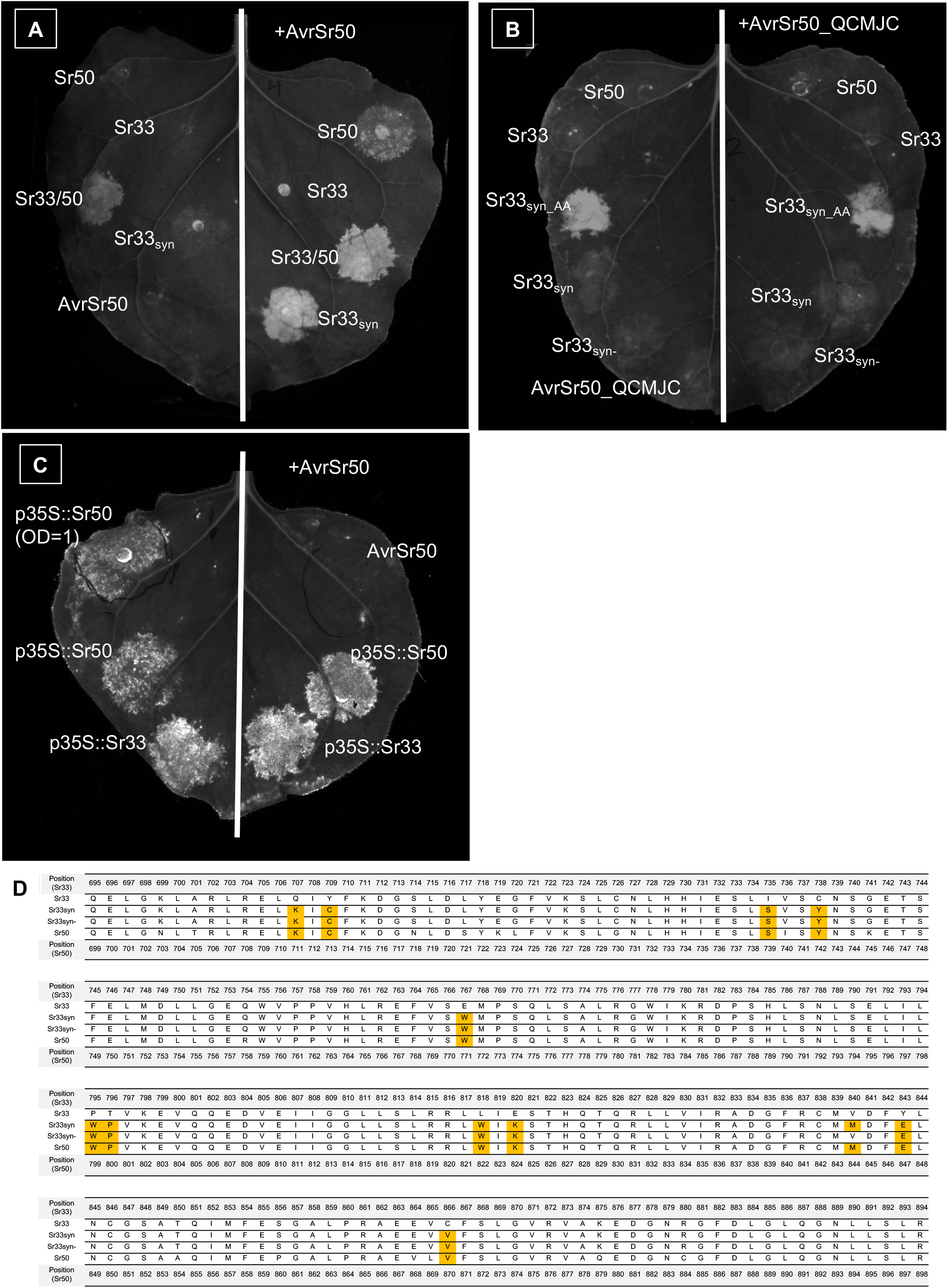
Pictures of *N. benthamiana* leaves of infiltrations of Sr33_syn_ and controls used to generate Figure 2C. Leaves of 4-5 weeks old *N. benthamiana* plants were infiltrated with *A. tumefaciens* carrying NLRs and effectors at OD_600_=0.3 and 0.4, respectively. UV images were taken on a BioRad Gel Imager 3-5 days after infiltration. **A.** Sr50, Sr33/50 and Sr33_syn_ induce cell death in response to AvrSr50 in *N. benthamiana*. **B.** None of the NLRs initiated cell death in response to the effector variant AvrSr50_QCMJC. **C.** Over-expression of Sr33 and Sr50 with *p35S* leads to cell death in absence of effector in *N. benthamiana*.

**Figure S4.**
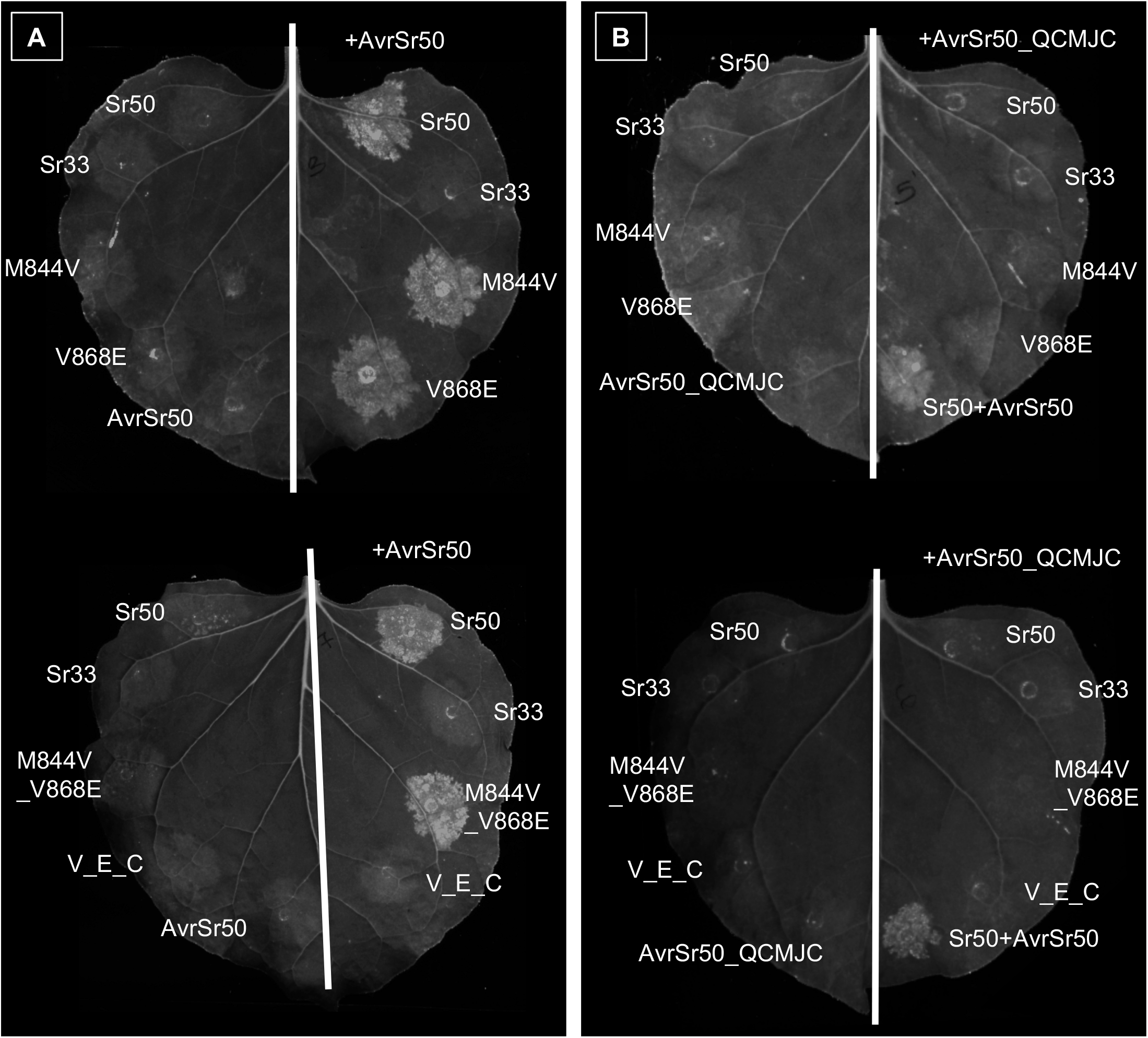
Pictures of *N. benthamiana* leaves infiltrated with Sr50 single and double mutants, and controls that were used to generate Figure 3B. Leaves of 4-5 weeks old *N. benthamiana* plants were infiltrated with *A. tumefaciens* carrying NLRs and effectors at OD_600_=0.3 and 0.4, respectively. UV images were taken on a BioRad Gel Imager 3-5 days after infiltration. **A.** Left panel. Sr50 wild type, single and double mutants induce cell death when co-infiltrated with AvrSr50. **B.** Right panel. Sr50 wild type, single and double mutants do not induce cell death when co-infiltrated with the effector variant AvrSr50_QCMJC.

**Figure S5.**
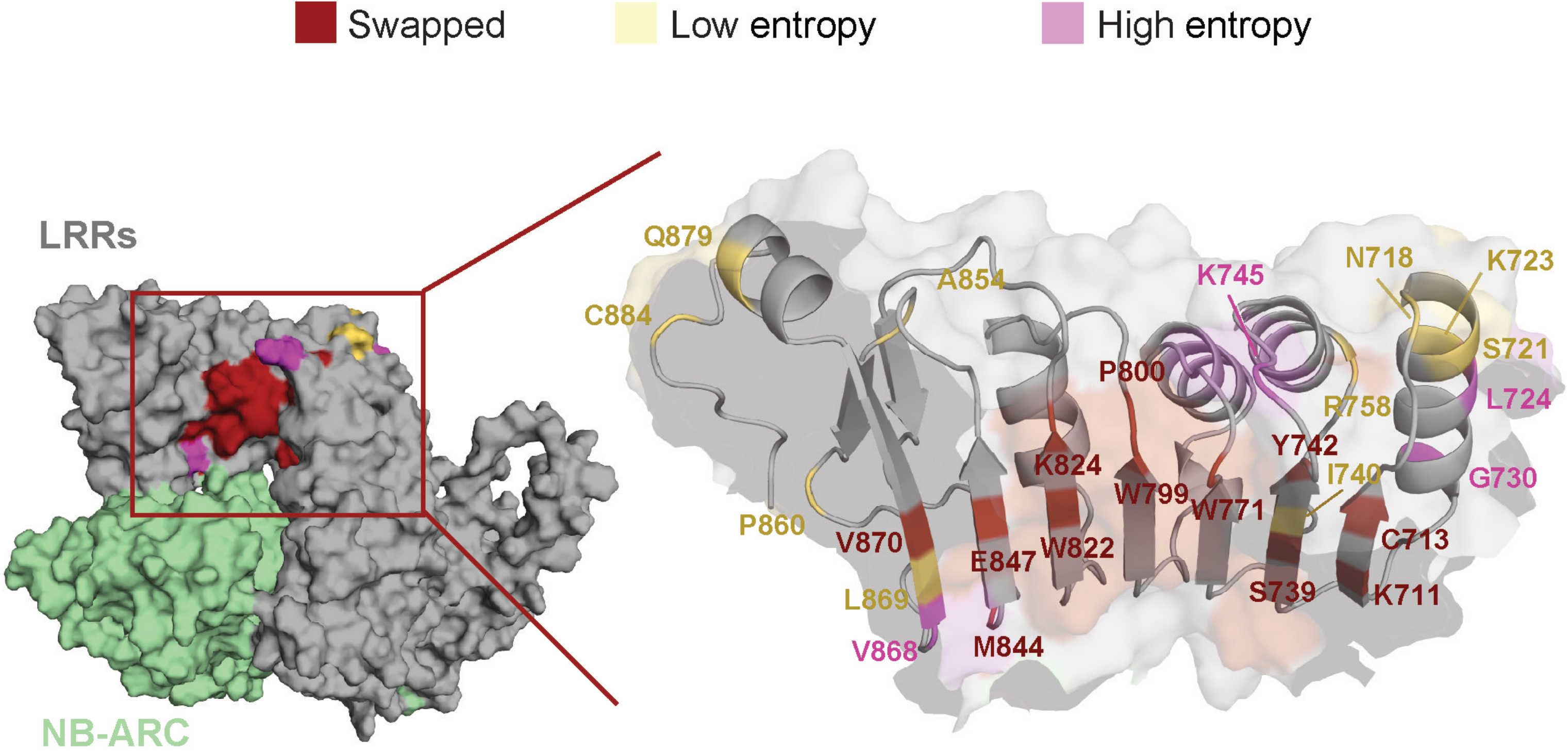
Variable LRR residues mapped into the Sr50 structure. The residues in the leucine-rich repeat (LRR) units 8-13 of Sr50 were compared to the LRR of Sr33. The sites that display polymorphism were mapped to the predicted protein structure of Sr50. The residues that display low and high entropy based on the entropy calculation of the Sr50 group were indicated in ivory and pink. The 12 residues used to synthesize Sr33_syn_ were colored in red and indicated as “swapped”.

**Figure S6.**
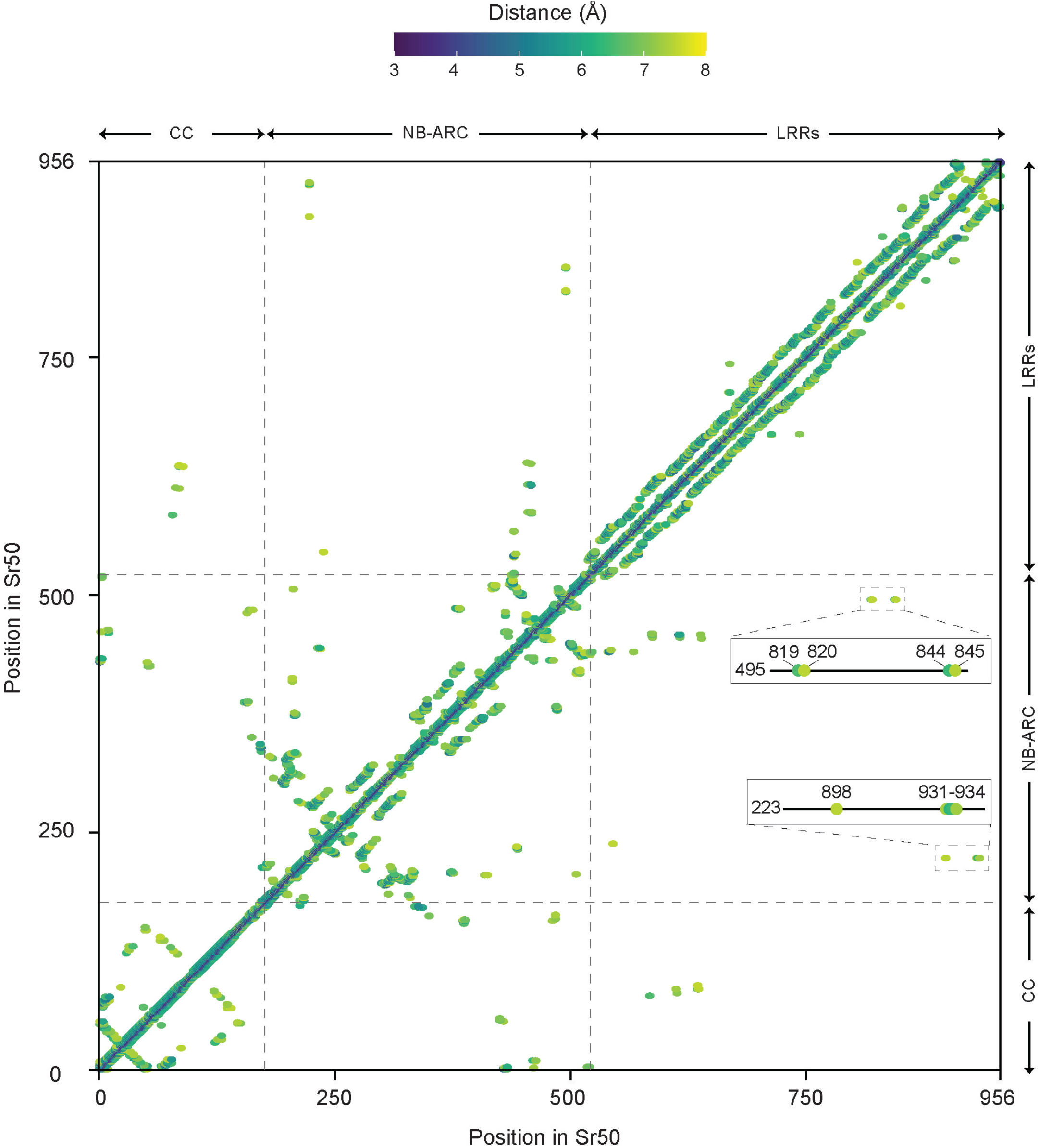
The inter-residual distances of Sr50. The pairwise distances between alpha carbon atoms in the predicted structure of Sr50 are depicted. The minimum observed distance was around 3Å. Only the pairs with the pairwise distance < 7.5Å were indicated. The x- and y-axis indicate amino acid position in Sr50. Multi-domain architecture (Coiled coil (CC), NB-ARC, and leucine-rich repeats (LRRs)) is provided around and across the axes. If a pair of residues are in proximity in the protein structure, a dot is indicated in their corresponding position. The insert captures pairwise distances between the residues in the NB-ARC domain and terminal LRRs.

